# Fast Lasso method for Large-scale and Ultrahigh-dimensional Cox Model with applications to UK Biobank

**DOI:** 10.1101/2020.01.20.913194

**Authors:** Ruilin Li, Christopher Chang, Johanne Marie Justesen, Yosuke Tanigawa, Junyang Qian, Trevor Hastie, Manuel A. Rivas, Robert Tibshirani

## Abstract

We develop a scalable and highly efficient algorithm to fit a Cox proportional hazard model by maximizing the *L*^1^-regularized (Lasso) partial likelihood function, based on the Batch Screening Iterative Lasso (BASIL) method developed in (Qian et al. 2019). The output of our algorithm is the full Lasso path, the parameter estimates at all predefined regularization parameters, as well as their validation accuracy measured using the concordance index (C-index) or the validation deviance. To demonstrate the effectiveness of our algorithm, we analyze a large genotype-survival time dataset across 306 disease outcomes from the UK Biobank (Sudlow et al. 2015). Our approach, which we refer to as snpnet-Cox, is implemented in a publicly available package.

## 1 Introduction

Survival analysis involves predicting time-to-event, such as survival time of a patient, from a set of features of the subject, as well as identifying features that are most relevant to time-to-event. Cox proportional hazard model (Cox 1972) provides a flexible mathematical framework to describe the relationship between the survival time and the features, allowing a time-dependent baseline hazard. Survival analysis faces computational and statistical challenges when the the predictors are ultrahigh-dimensional (when feature dimension is greater than the number of observations) and large scale (when the data matrix does not fit in the memory). Based on the Batch Screening Iterative Lasso (BASIL), we develop an algorithm to fit a Cox proportional hazard model by maximizing the Lasso partial likelihood function. We apply the method to 306 time-to-event disease outcomes from UK Biobank combined with genetic data. We generate improved predictive models with sparse solutions using genetic data with the number of variables selected ranging from a single active variable in the set and others with almost 2,000 active variables.

### 1.1 Cox Proportional Hazard Model

Given a numerical predictor *X* ∈ ℝ^*d*^, Cox model assumes that there exists a baseline hazard function *h*_0_ : ℝ^+^ ↦ ℝ^+^ and a parameter vector *β* ∈ ℝ^*d*^ such that the hazard function for survival time has the form:

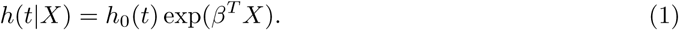

Intuitively the hazard function at time *t* measures the relative risk of death around time *t*, given that the patient survives up to time *t*. Under Cox proportional hazard model, the hazard ratio between two subject with covariates *X* and *X*′ can be written as:

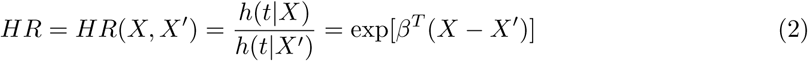

When *X* is an indicator for a treatment, the hazard ratio can be interpreted as the risk of event occurring in the treatment group, compared to the risk in the control group, and the regression coefficient *β* is the log-hazard ratio.

To describe the distribution of the survival time we can equivalently use its cumulative distribution function:

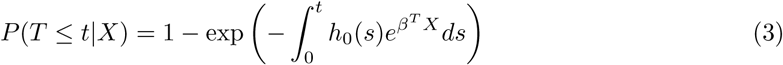

In practice it is often the case that the survival time is right-censored. That is the event has not yet happened at the time the data was collected. Therefore for the *i*th individual we observe a tuple (*X*_*i*_, *O*_*i*_, *T*_*i*_), where *X*_*i*_ ∈ ℝ^*d*^ is the predictors, *O*_*i*_ ∈ {0, 1} is the event indicator. If *O*_*i*_ = 1, then *T*_*i*_ is the actual survival time of the *i*th individual. If *O*_*i*_ = 0, then we only know that the true survival time of the *i*th individual is longer than *T*_*i*_. Throughout this paper we will assume that the censoring is non-informative, meaning that the time of censoring is independent of the (possibly unobserved) event time conditional on *X*_*i*_.

One advantage of the Cox model is that, while being a semi-parametric model (the baseline function is non-parametric), we could still estimate the parameter *β* without estimating the baseline function. This can be achieved by choosing 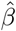 that maximizes the log-partial likelihood function:

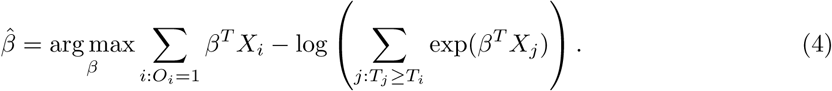

We use the C-index to evaluate a fitted 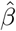:

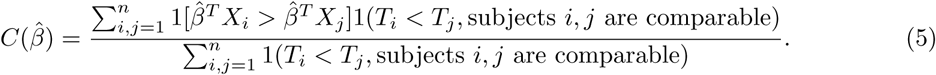

Here subject *i* and *j* are comparable if and only if:

- *O*_*i*_ = *O*_*j*_ = 1. That is an event has been observed for both *i* and *j*;
- *O*_*i*_ = 0, *O*_*j*_ = 1, *T*_*i*_ > *T*_*j*_, or vice-versa. In this case, although only one event event is observed, the ordering of the underlying event time can still be inferred.

In this paper we assume that there is no ties between event time or predictors. For a more complete description of C-index, see (Harrell et al. 1982, Li & Tibshirani 2019)

## 2 Method

### 2.1 Preliminaries

We first introduce the following notations:

- Let *n, d* be the number of observations and the number of features respectively. Let ***X*** ∈ ℝ^*n*×*d*^ be the matrix of predictors. To simplify notation, we use *n, d*, ***X*** for all of train, test, and validation set. Whether ***X*** comes from train, test, or validation data can be inferred from the context.
- Let *X*_*i*_ ∈ ℝ^*d*^ be the *i*th row of ***X***.
- Let *x*_*j*_ ∈ ℝ^*n*^ be the *j*th column of ***X***.
- Denote the log-partial likelihood function as *f* (*β*). That is

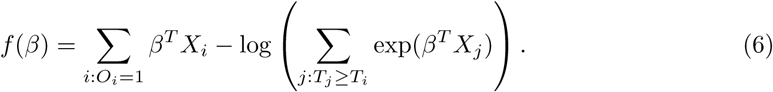

We focus on survival analysis in the high-dimensional regime, where the number of predictors is greater than the number of observations (*d* > *n*), although same procedure can easily be applied to low-dimensional cases. We use Lasso to perform variable selection and estimation at the same time. In particular, we optimize the *L*^1^-regularized log-partial likelihood:

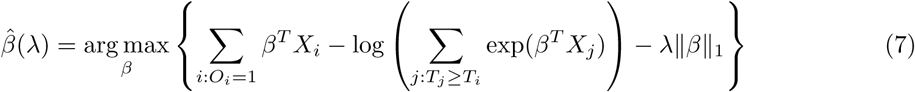

where the 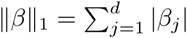. More generally, we allow each parameter or each observation to have a different weight in the objective function, the right-hand side of (7). In particular, given a vector of penalty factors 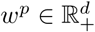, and observation weight 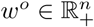, we define the general objective function to be

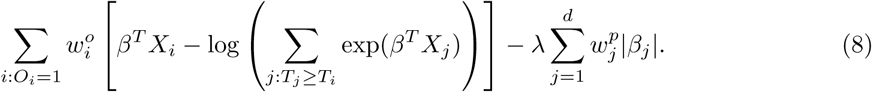

This can be particularly useful if we are considering genetic variants that we would like to upweight during variable selection, e.g. coding variants in a region of perfect linkage disequilibrium. To simplify the notation we describe our algorithm assuming that the parameters and the observations are unweighted.

### 2.2 Hyperparameter Selection

To find the optimal hyperparameter *λ*, we start with a sequence of *L* candidate regularization parameters *λ*_1_ > *λ*_2_ > …, *λ*_*L*_ > 0 and compute the corresponding parameter estimate as well as the validation metric. The optimal regularization parameter is then selected to be *λ*_*l*_ that maximizes the validation metric and 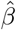 is set to be 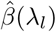. The sequence of regularization parameters can be chosen by setting *λ*_1_ to be sufficiently large such that the optimal 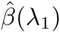 is just zero, and find *L* = 100 equally spaced *λ*s in log-scale.

Applying this procedure naively requires solving *L* optimization problems, each reading the entire predictor matrix ***X***. The key components of our algorithm that significantly speed up the computation are the following observations adapted from (Qian et al. 2019).

### 2.3 Batching Screening Procedure

The KKT condition of (7) indicates that the optimal 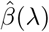 must satisfy:

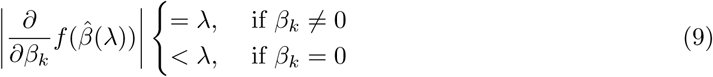

When *λ* is sufficiently large, 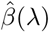 is sparse, so our strategy is to solve the optimization problem (7) using only a small subset of features, assuming all the others have coefficient zero. Then we verify that the solution satisfies the KKT condition (9). We iteratively apply this strategy for *λ* = *λ*_1_, …, *λ*_*L*_ to obtain the entire Lasso path. To determine which predictors to include in the model, we adopt the screening rules used in BASIL, which is inspiered by the strong rules proposed in (Tibshirani et al. 2012). In Cox model, the strong rules assumes 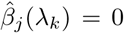 (discard the *j*th predictor when fitting) if

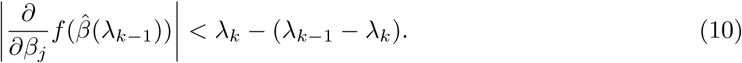

By convention we set *λ*_0_ =∞. Although it is possible for strong rules to fail, it rarely happens when *d* > *n*.

Before we describe the full algorithm, first we write the gradient of the log-partial likelihood into a simple form. Notice that the gradient of the log-partial likelihood function can be written as:

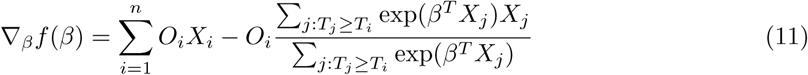

Let *r* = *r*(*data, β*) ∈ ℝ^*n*^ be defined as

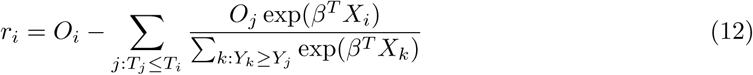

Then by direct computation one can show that

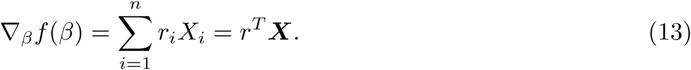

Our full algorithm follows the same structure as in BASIL (Qian et al. 2019), where at each iteration of our algorithm we look for Lasso solution for multiple consecutive *λ*s in the Lasso path so that large dataset is not read in to frequently. Suppose 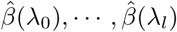 have been computed in the first *k* − 1 iterations. At the *k*th iteration we maintain a strong set 𝒮^(*k*)^ ⊆ [*d*], an ever-active set 𝒜^(*k*)^ ⊆ [*d*], and a set of regularization parameters Λ^(*k*)^ ⊆ {*λ*_1_, …, *λ*_*L*_} that we use to fit Lasso in the current iteration. Here 𝒮^(*k*)^ is the small subset of variables that are used to fit 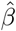 implied by our screening procedure (10), and 𝒜^(*k*)^ is the subset of variables such that Lasso coefficients are non-zero for at least one *λ* in *λ*_0_,, *λ*_*l*_. Each iteration has the following three steps: screening, fitting and checking. At the screening step we update ^(*k*)^, which includes variables that are in 𝒜^(*k*−1)^ and the top *M* variables with the largest partial derivative in the left-hand side of (10) that are also not in 𝒜^(*k*−1)^. At the fitting step, we solve the optimization problem (7) using regularization parameters in Λ^(*k*)^ and variables in 𝒮^(*k*)^. At the checking step, we check that the KKT condition (9) is satisfied and extend Λ^(*k*)^ to Λ^(*k*+1)^ by adding the next few *λ*s in the list that are unused yet. As an optional step, we compute the validation C-index of 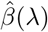 for *λ* ∈ Λ^(*k*)^. Algorithm 1 summarizes this procedure.

#### Algorithm 1: BASIL for Cox Model

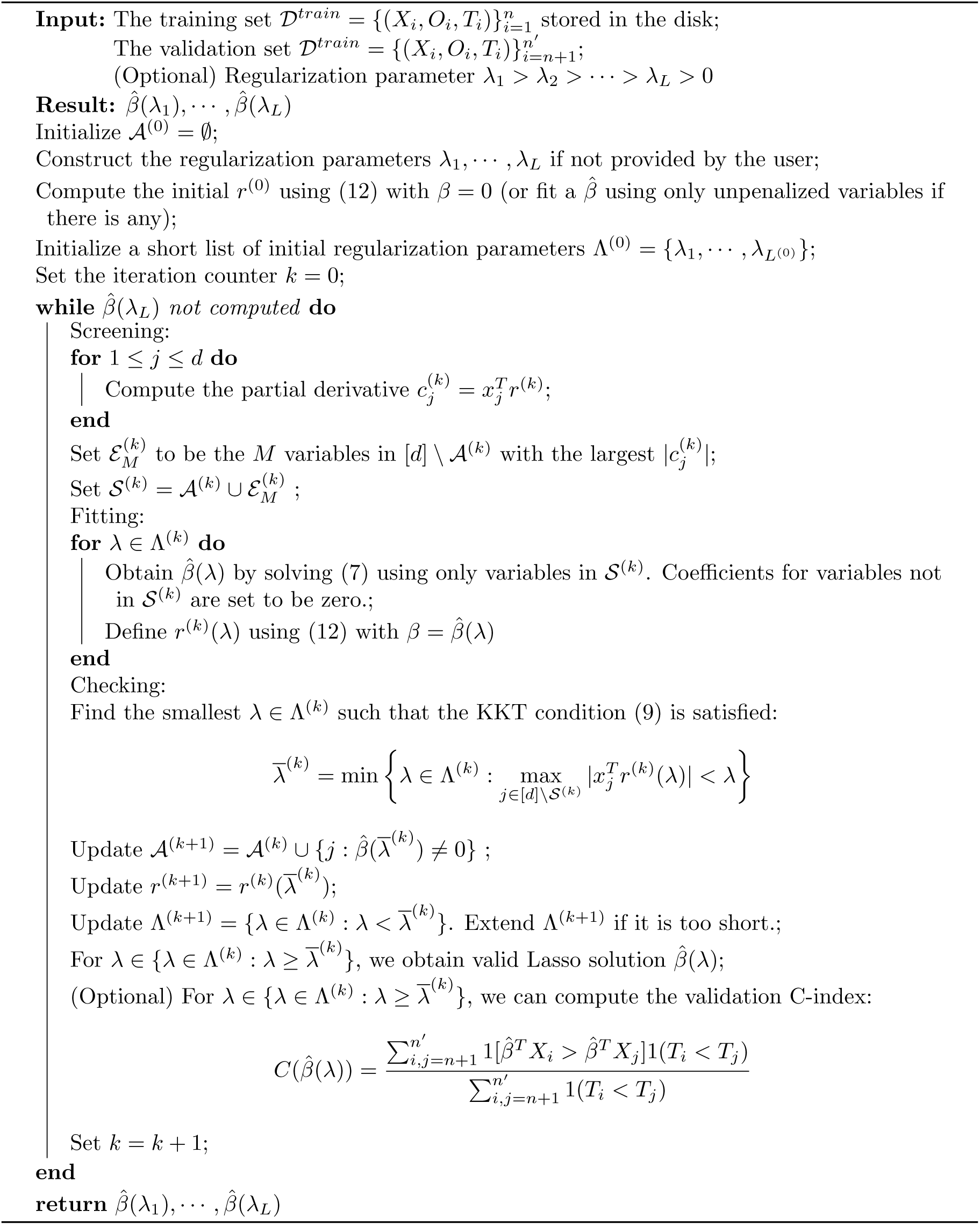

### 2.4 Fast C-Index Computation

Several frequently-used C-index computational algorithms, including the first algorithm we tried, have time complexity 𝒪(*n*^2^). As population-scale cohorts, like UK Biobank, Million Veterans Program, and FinnGen, aggregate time to event data for survival analysis it is increasingly important to consider the computational costs of statistics like C-index to build and evaluate predictive models. The time to event data for survival analysis include age of disease onset, progression from disease diagnosis to another more severe outcome, like surgery or death. Here, we present an implementation with 𝒪(*n* log *n*) time complexity (and 𝒪(*n*) space complexity) that can introduce over 10,000x speedup for biobank-scale data relative to several R packages, and over 10x speedup compared to existing 𝒪(*n* log *n*) tim complexity (and 𝒪(*n* + *n* log *n*) space complexity) algorithm implemented in the survival analysis package (Therneau & Lumley 2014).

We initially assume there are no tied *f* or *T* values.

I. Define *g* (*X*_*j*_) to be the number of values *i* in {1, …, *n*} where 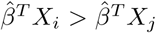. For example, if *n* = 3, *f* (*X*_1_) = 6, *f* (*X*_2_) = 3, and *f* (*X*_3_) = 4, then *g* (*X*_1_) = 2, *g* (*X*_2_) = 0, and *g* (*X*_3_) = 1. Note that 1 [*g* (*X*_*i*_) < *g* (*X*_*j*_)] is always the same as 1 [*f* (*X*_*i*_) < *f* (*X*_*j*_)]; these functions have the same rank ordering. So we just work with *g* in the remainder of this discussion. And, given *f*, it is straightforward to compute *g* in 𝒪(*n* log *n*) time.
II. Sort the records in order of nonincreasing *T*. This lets us rewrite 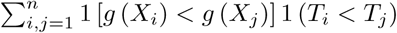 as 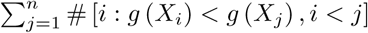 for uncensored *j*.
III. Each *g* (*X*) value is a distinct nonnegative integer in {0, 1, …, *n* − 1}. So we can use a bitarray to represent the set {*g* (*X*_*i*_) : *i* < *j*}. This bitarray has two nice properties:
  a. # [*i* : *g* (*X*_*i*_) < *g* (*X*_*j*_), *i* < *j*] is the number of set bits (“popcount”) in bitarray positions {0, 1, …, *g* (*X*_*j*_) − 1}.
  b. When we advance to the next *j*, we set one bit in the bitarray to update it. Bitarrays are compact, and very efficient to work with. (The exact arithmetic and bitwise operations we used were primarily informed by (Knuth 2011) and (Mula et al. 2016).) However, we need to perform 𝒪(*n*) array-popcount operations, so the top-level algorithm is still 𝒪(*n*^2^) if each popcount takes 𝒪(*n*) time. To get the array-popcount operations down to 𝒪(log *n*), we augment the bitarray with a stack of indexes. For example, the first-level index can be of the form <# of set bits in {0, 1, …, 511}>, <# of set bits in {512, 513, …, 1023}>,<# of set bits in {1024, 1025, …, 1535}>,…, and the second-level index can be of the form <# of set bits in {0, 1, …, 16383}>,<# of set bits in {16384, 16385, …, 32767}>, … Then, for *g*(*X*_*j*_) = 40000, we wouldn’t actually need to scan the first 40000 bits. We could instead get popcount({0, 1, …, 39935}) by adding 2 second-level-index entries to 14 first-level-index entries, and we’d only need to directly scan the last 64 bits. The drawback is that when we advance to the next *j*, we now have to update the indexes in addition to setting a bit in the main bitarray. But in the example above, only two index entries need to be updated. In the general case, the number of required index updates when we increment *j* is 𝒪(log *n*), which is within our 𝒪(*n* log *n*) budget.
IV. Ties introduce some challenges to deal with, but the same core algorithm is still effective. With very large groups of tied *f* () values, the main tie-handling strategy causes the algorithm to degrade to 𝒪(*n*^2^), so our implementation detects this scenario and switches to a slightly different loop that doesn’t degrade.

## 3 Applications

### 3.1 UK Biobank age of diagnosis data preparation

We have prepared an age of diagnosis dataset from the UK Biobank derived from Category 1712, the category containing data showing the ‘first occurrence’ of any code mapped to 3-character ICD-10 (see Supplementary Material).

Briefly, the data-fields have been generated by mapping: Read code information in the Primary Care data (Category 3000); ICD-9 and ICD-10 codes in the Hospital inpatient data (Category 2000); ICD-10 codes in Death Register records (Field 40001, Field 40002), and Self-reported medical condition codes (Field 20002) reported at the baseline or subsequent UK Biobank assessment centre visit to 3-character ICD-10 codes.

For each code two data-fields are available: the date the code was first recorded across any of the sources listed above, the source where the code was first recorded, and information on whether the code was recorded in at least one other source subsequently.

We used these data and computed an age of diagnosis by using the Month of Birth Data Field (Data-Field 52) and Year of Birth (Data-Field 34).

### 3.2 Genetic data preparation

Here, we used genotype data from the UK Biobank dataset release version 2 and the hg19 human genome reference for all analyses in the study. To minimize the variabilities due to population structure in our dataset, we restricted our analyses to include 337, 151 unrelated White British individuals, used sex, Array (UK Biobank was genotyped in two different platforms), and 10 principal components derived from the genotype data as covariates (described in detail in Supplementary Note).

We focused our analysis on variants with a minor allele frequency (MAF) greater than or equal to 0.1% for directly genotyped variants in either array, in addition to the Human Leukocyte Antigen (HLA) alleles (Bycroft et al. 2018) and copy number variants (CNVs) described in (Aguirre et al. 2019) for a total of 1.08 million variants.

We split our dataset into a 70% training (n = 236,004), 10% validation, (n = 33,716) and 20% held out test set (n = 67,430), and apply snpnet-Cox with 50 iterations. We focus our analysis on 306 ICD10 codes with at least 950 cases in the 337, 151 individuals dataset.

### 3.3 snpnet-Cox results

We summarize the results across the 306 ICD10 codes, but focus our detailed analysis for four of them including:

1. asthma (ICD10 code: J45),
2. gout (M10),
3. disorders of porphyrin and bilirubin metabolism (E80), and
4. atrial fibrillation and flutter (I48).

When assessing the predictive performance of 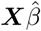, where 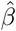 is the fitted regression coefficients from the snpnet-Cox, and ***X*** is the matrix of genotypes for the individuals in the held out test set for variants with non-zero regression coefficients from snpnet-Cox, we applied Cox proportional hazard model in the survival package. We applied a couple of procedures to give a high level overview of the results. First, we assessed whether the Polygenic Hazard Score (PHS), or 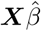, was significantly associated to the time to event data in the held out test set (so that we obtained a *P*-value for each ICD10 code). Second, we computed the Hazards Ratio (HR) for the scale (standard deviation unit), and different thresholded percentiles (top 1%, 5%, 10%, and bottom 10% compared to the 40-60%) of 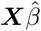. Third, we computed the C-index (Harrell et al. 1982).

The C-index for the 101 ICD10 codes with PHS *P* < 0.01 range from 0.511 to 0.884 (see Global Biobank Engine (GBE) snpnet-Cox page https://biobankengine.stanford.edu/snpnetcox) and HR per standard deviation of PHS from 1.042 to 13.167. The results further highlight the sparsity property of Lasso in the Cox model implemented in snpnet-Cox with some ICD10 codes including a single active variable in the set and others with almost 2,000 active variables (e.g. non-insulin-dependent diabetes mellitus).

#### 3.3.1 Asthma - J45

Motivated by the varying age of asthma onset, a common disease that affects a substantial fraction of young adults, we hypothesized that a PHS could capture individuals that are not only at higher risk of disease onset, but also at a higher risk of developing asthma at a younger age.

Here, we estimate a HR of 1.428 per SD of PHS (C-index of 0.605), and HR of 2.740, 2.137 and 1.825 for the top 1, 5, and 10% of the PHS distribution compared to the 40 − 60%. Further, we find that 14.2% of individuals in the top 1% of the PHS score developed asthma by age 20.5 compared to only 1.1% in the bottom 10% and 3.2% of the 40 − 60%ile of the PHS score (see Figure 1), which underscores the relevance of PHS in the context of early onset of common diseases that are hypothesized to have a monogenic signature (Kelsen & Baldassano 2017). The asthma PHS is composed of 1.567 active variable of which some are known from previous GWAS of traits related to asthma. As an example, we identify the rs2381416 (MAF = 0.26) upstream of *GTF3AP1* to associate with asthma with an effect size of -0.11. This variant has previously been found to associate with eosinophil count (Gudbjartsson et al. 2009) and severity of childhood asthma (Smith et al. 2017).

**Figure 1:**
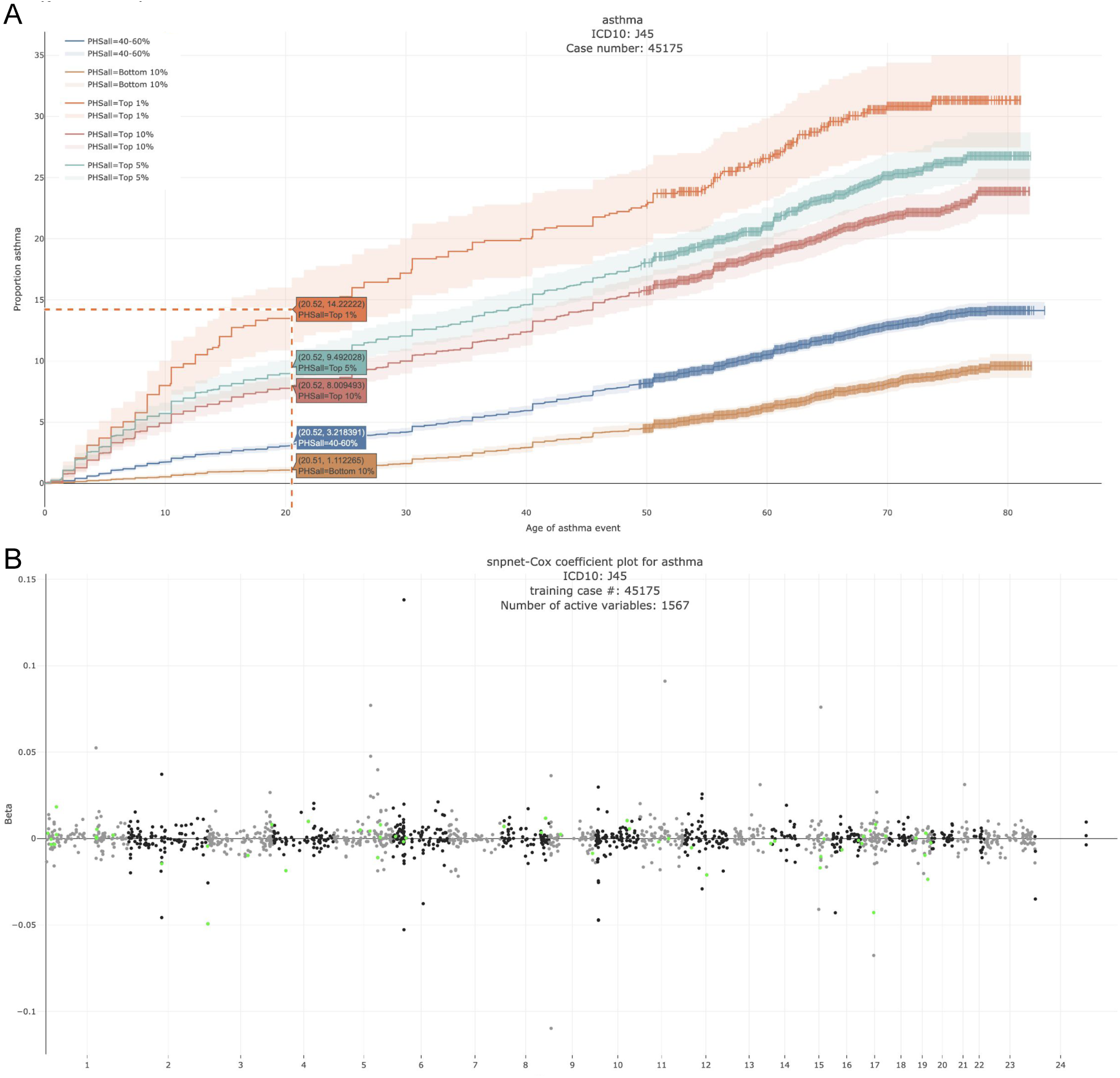
Asthma. A. Kaplan-Meier curves for percentiles of Polygenic Hazard Scores (PHS), or 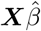, for variants selected by snpnet-Cox, in the held out test set (orange - top 1%, green - top 5%, red - top 10%, blue - 40-60%, and brown - bottom 10%; ticks represent censored observations. Highlighted are the proportion of asthma events by age 20 across the percentile scores. B. Plot of snpnet-Cox coefficients for asthma with 1, 567 active variables. Green dots represent protein-altering variants.

#### 3.3.2 Gout - M10

Gout is a common disease, affecting at least 1% of men in Western countries, with a strong male to female imbalance (Terkeltaub 2003). It is a form of arthritis caused by excess uric acid in the bloodstream and characterized by severe pain, redness, and tenderness in joints.

In the UK Biobank study, we estimate a Hazard Ratio of 1.679 per SD of PHS (C-index of 0.649), and HR of 3.70, 2.502 and 2.073 for the top 1,5, and 10% of the PHS distribution compared to the 40 − 60%. Further, we find that 4.89% of individuals in the top 1% of the PHS score developed asthma by age 50.1 compared to only 0.30% in the bottom 10% and 1.02% of the 40 − 60%ile of the PHS score (see Figure 2). The gout PHS consists of 1.970 active variables and we identify loci that have been identified in prior GWAS (Dehghan et al. 2008).

**Figure 2:**
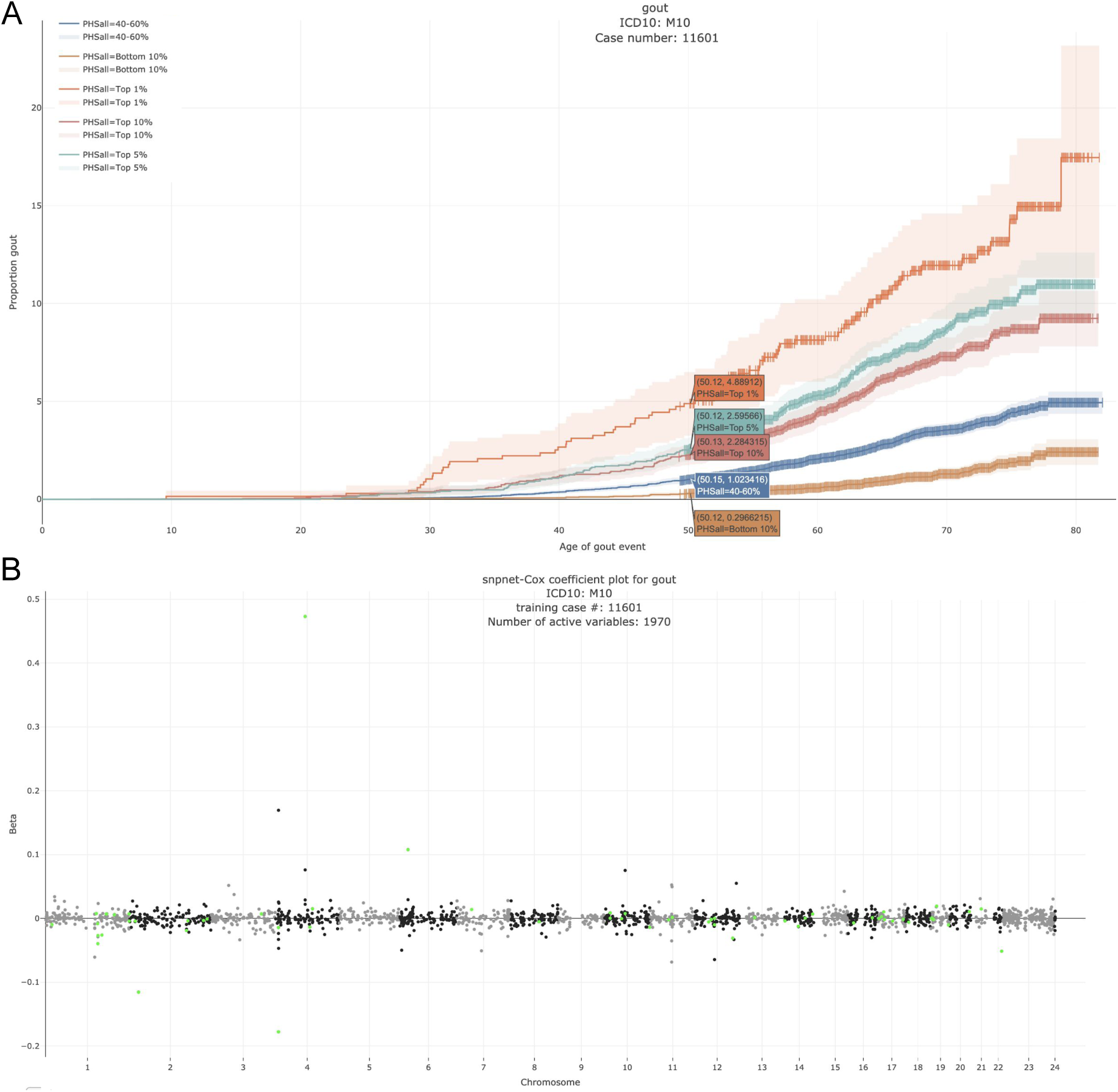
Gout. A. Kaplan-Meier curves for percentiles of Polygenic Hazard Scores (PHS), or **X**^T^ **B**, for variants selected by snpnet-Cox, in the held out test set (orange - top 1%, green - top 5%, red - top 10%, blue - 40-60%, and brown - bottom 10%; ticks represent censored observations. Highlighted are the proportion of gout events by age 50 across the percentile scores. B. Plot of snpnet-Cox coefficients for gout with 1, 970 active variables. Green dots represent protein-altering variants.

#### 3.3.3 Disorders of porphyrin and bilirubin metabolism - E80

Bilirubin, which is the principal component of bile pigments, is the end product of the catabolism of the heme moiety of hemoglobin and other hemoproteins. If bilirubin is produced in excessive amounts or hepatic excretion of bilirubin into bile is defective, the concentration of bilirubin in the blood and tissues increases, which may result in jaundice (Bosma 2003), a well recognisable symptom of liver disease.

We estimate a HR of 13.167 per SD of PHS (C-index of 0.884). Here, given that we have only 2 active variables, we find that the snpnet-Cox algorithm finds a sparse solution (see Figure 3). One of the active variables is the intron variant (rs6742078) of *UTG1A4* (MAF = 0.31) which encodes an enzyme UDP-glucuronosyltransferase that transforms small lipophilic molecules such as bilirubin (Tukey & Strassburg 2000).

**Figure 3:**
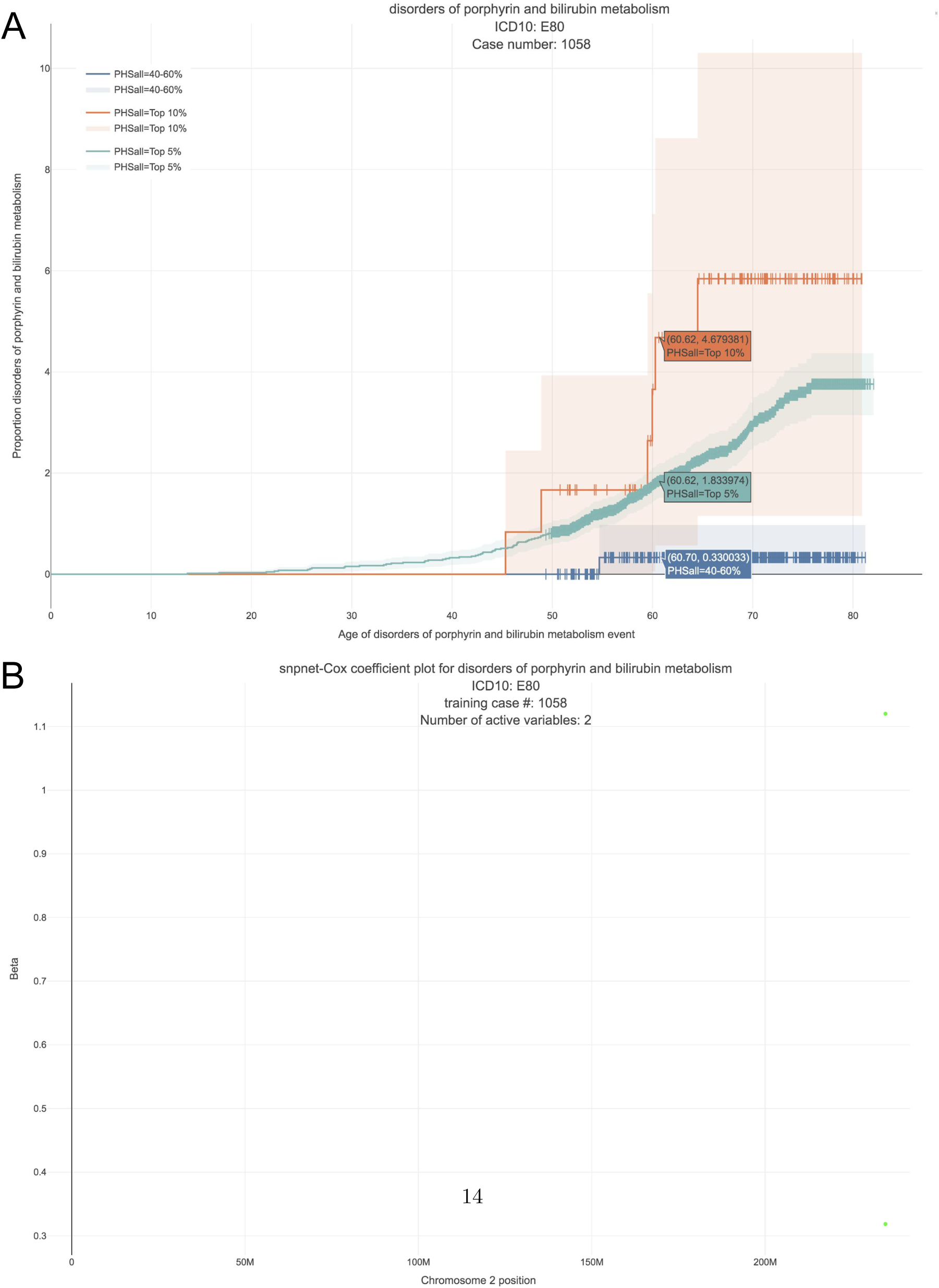
Disorders of porphyrin and bilirubin metabolism. A. Kaplan-Meier curves for percentiles of Polygenic Hazard Scores (PHS), or **X**^T^ **B**, for variants selected by snpnet-Cox, in the held out test set (orange - top 1%, green - top 5%, red - top 10%, blue - 40-60%, and brown - bottom 10%; ticks represent censored observations. Highlighted are the proportion of disorders of porphyrin and bilirubin metabolism events by age 60 across the percentile scores. B. Plot of snpnet-Cox coefficients for disorders of porphyrin and bilirubin metabolism with 2 active variables. Green dots represent protein-altering variants.

#### 3.3.4 Atrial fibrillation and flutter - I48

Atrial fibrillation is the most common type of arrhythmia in adults. The prevalence increases from less than 1% in persons younger than 60 years of age to more than 8% in those older than 80 years of age (McNamara et al. 2003). Earlier onset of atrial fibrillation is believed to have a strong genetic component and whether that has more of a polygenic or monogenic flavor is currently unknown.

In the UK Biobank study we estimate a Hazard Ratio of 1.466 per SD of PHS (C-index of 0.618), and HR of 3.883, 2.319 and 1.861 for the top 1,5, and 10% of the PHS distribution compared to the 40 − 60%. Further, we find that 6.57% of individuals in the top 1% of the PHS score developed asthma by age 60 compared to only 0.70% in the bottom 10% and 1.41% of the 40 − 60%ile of the PHS score (see Figure 4), which underscores the relevance of PHS in the context of early onset of atrial fibrillation.

**Figure 4:**
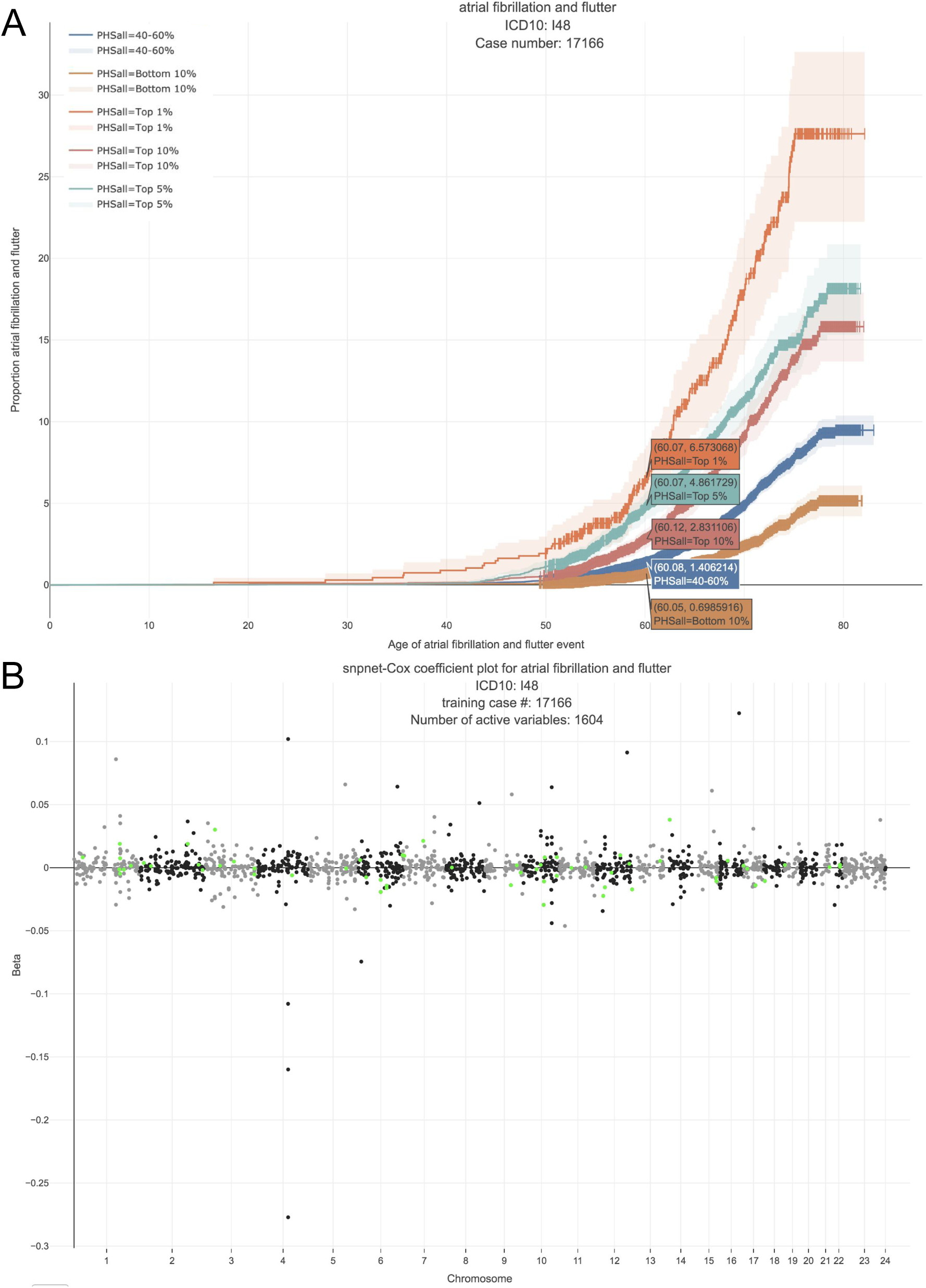
Atrial fibrillation. A. Kaplan-Meier curves for percentiles of Polygenic Hazard Scores (PHS), or **X**^T^ **B**, for variants selected by snpnet-Cox, in the held out test set (orange - top 1%, green - top 5%, red - top 10%, blue - 40-60%, and brown - bottom 10%; ticks represent censored observations. Highlighted are the proportion of atrial fibrillation events by age 60 across the percentile scores. B. Plot of snpnet-Cox coefficients for atrial fibrillation with 1, 604 active variables. Green dots represent protein-altering variants.

**Figure 5:**
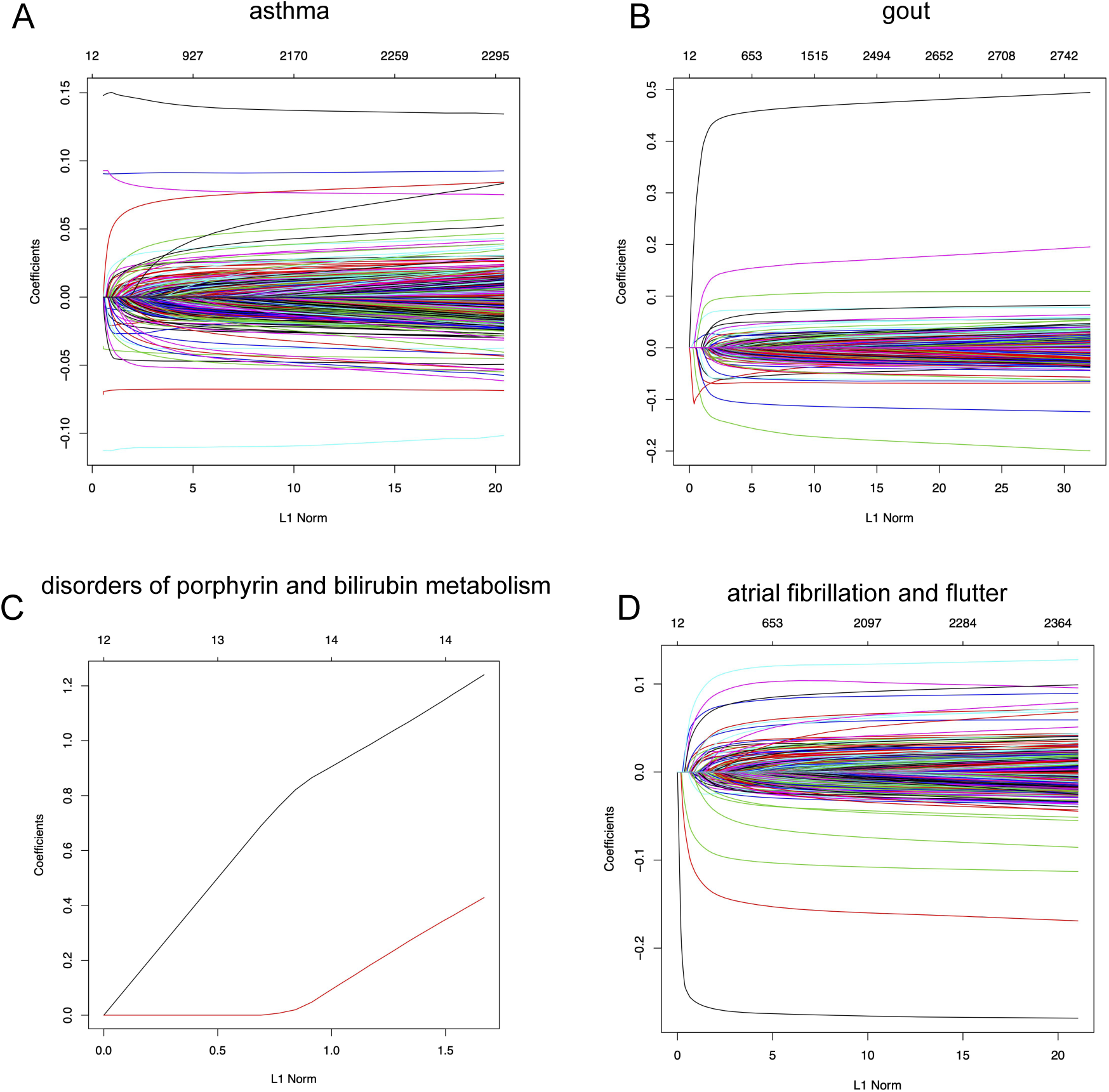
snpnet-Cox paths. Each line in these plots corresponds to a variable from the best model. The vertical axis represents the L_1_ norm of the estimated coefficients and the horizontal axis represents the value of the coefficients. The path is computed at various level of regularization parameter. The whiskers at the top of the plot are the number of variables selected. The first 12 variables are the covariates including age, sex, PC1-10.

## 4 Discussion

In this article, we developed the batch screening iterative LASSO (BASIL) algorithm (Qian et al. 2019) to find the lasso path of Cox proportional hazard models. We implemented an optimized C-index function, which computes the C-index of a fitted Cox model in *O*(*n* log *n*) time with an excellent constant factor. Our method was applied to the UK Biobank dataset to identify genetic variants that are associated with time-to-event phenotypes and to build Polygenic Hazard Scores (PHS). Visualizations of snpnet-Cox results across 306 ICD10 codes are available in Global Biobank Engine (https://biobankengine.stanford.edu/snpnetcox) (McInnes et al. 2018).

Our current approach does have limitations, which we hope to resolve in future work. First, we assume that each individual has independent survival times (conditional on the features). This may become a limitation as population-scale cohorts especially in population isolates like in Finland sample related individuals. Second, we do not provide procedures for false discovery rate estimates based on selected features, which may be useful in communicating confidence in a single active variable (Taylor & Tibshirani 2015). Third, as we move towards whole genome sequencing data where a large fraction of variants discovered will have a rare event property, i.e. observed in a handful of individuals, the validation accuracy may need to be redefined to evaluate a fitted 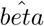. Fourth, we do not consider time-varying coefficients and time-varying covariates, which may improve inference in the setting where features may have multiple measurements over time. These are areas of future direction that we anticipate we will address.

We provide the implementation of our approach in a publicly available package `snpnet` available at https://github.com/junyangq/snpnet with `cindex` package dependency available at https://github.com/chrchang/plink-ng/tree/master/2.0/cindex.

## References

Aguirre, M., Rivas, M. & Priest, J. (2019), ‘Phenome-wide burden of copy number variation in uk biobank’, American Journal of Human Genetics pp. 373–383.

Bosma, P. J. (2003), ‘Inherited disorders of bilirubin metabolism’, Journal of hepatology 38(1), 107–117.

Bycroft, C., Freeman, C., Petkova, D., Band, G., Elliott, L. T., Sharp, K., Motyer, A., Vukcevic, D., Delaneau, O., O’Connell, J. et al. (2018), ‘The uk biobank resource with deep phenotyping and genomic data’, Nature 562(7726), 203.

Cox, D. R. (1972), ‘Regression models and life-tables’, Journal of the Royal Statistical Society. Series B (Methodological) 34(2), 187–220. URL: http://www.jstor.org/stable/2985181

Dehghan, A., Köttgen, A., Yang, Q., Hwang, S.-J., Kao, W. L., Rivadeneira, F., Boerwinkle, E., Levy, D., Hofman, A., Astor, B. C. et al. (2008), ‘Association of three genetic loci with uric acid concentration and risk of gout: a genome-wide association study’, The Lancet 372(9654), 1953–1961.

Gudbjartsson, D. F., Bjornsdottir, U. S., Halapi, E., Helgadottir, A., Sulem, P., Jonsdottir, G. M., Thorleifsson, G., Helgadottir, H., Steinthorsdottir, V., Stefansson, H. et al. (2009), ‘Sequence variants affecting eosinophil numbers associate with asthma and myocardial infarction’, Nature genetics 41(3), 342.

Harrell, F. E., Califf, R. M., Pryor, D. B., Lee, K. L. & Rosati, R. A. (1982), ‘Evaluating the yield of medical tests’, Jama 247(18), 2543–2546.

Kelsen, J. R. & Baldassano, R. N. (2017), ‘The role of monogenic disease in children with very early onset inflammatory bowel disease.’, Current opinion in pediatrics 29(5), 566–571.

Knuth, D. E. (2011), The art of computer programming, volume 4A: combinatorial algorithms, part 1, Pearson Education India.

Li, R. & Tibshirani, R. (2019), ‘On the use of c-index for stratified and cross-validated cox model’, arXiv preprint 1911.09638.

McInnes, G., Tanigawa, Y., DeBoever, C., Lavertu, A., Olivieri, J. E., Aguirre, M. & Rivas, M. (2018), ‘Global biobank engine: enabling genotype-phenotype browsing for biobank summary statistics’, BioRxiv p. 304188.

McNamara, R. L., Tamariz, L. J., Segal, J. B. & Bass, E. B. (2003), ‘Management of atrial fibrillation: review of the evidence for the role of pharmacologic therapy, electrical cardioversion, and echocardiography’, Annals of internal medicine 139(12), 1018–1033.

Mu la, W., Kurz, N. & Lemire, D. (2016), ‘Faster population counts using avx2 instructions’, arXiv preprint 1611.07612.

Qian, J., Du, W., Tanigawa, Y., Aguirre, M., Tibshirani, R., Rivas, M. A. & Hastie, T. (2019), ‘A fast and flexible algorithm for solving the lasso in large-scale and ultrahigh-dimensional problems’, bioRxiv. URL: https://www.biorxiv.org/content/early/2019/05/07/630079

Smith, D., Helgason, H., Sulem, P., Bjornsdottir, U. S., Lim, A. C., Sveinbjornsson, G., Hasegawa, H., Brown, M., Ketchem, R. R., Gavala, M. et al. (2017), ‘A rare il33 loss-of-function mutation reduces blood eosinophil counts and protects from asthma’, PLoS genetics 13(3), e1006659.

Sudlow, C., Gallacher, J., Allen, N., Beral, V., Burton, P., Danesh, J., Downey, P., Elliott, P., Green, J., Landray, M., Liu, B., Matthews, P., Ong, G., Pell, J., Silman, A., Young, A., Sprosen, T., Peakman, T. & Collins, R. (2015), ‘Uk biobank: An open access resource for identifying the causes of a wide range of complex diseases of middle and old age’, PLOS Medicine 12(3), 1–10. URL: https://doi.org/10.1371/journal.pmed.1001779

Taylor, J. & Tibshirani, R. J. (2015), ‘Statistical learning and selective inference’, Proceedings of the National Academy of Sciences 112(25), 7629–7634.

Terkeltaub, R. A. (2003), ‘Gout’, New England Journal of Medicine 349(17), 1647–1655.

Therneau, T. M. & Lumley, T. (2014), ‘Package ‘survival”, Survival analysis Published on CRAN.

Tibshirani, R., Bien, J., Friedman, J., Hastie, T., Simon, N., Taylor, J. & Tibshirani, R. J. (2012), ‘Strong rules for discarding predictors in lasso-type problems’, Journal of the Royal Statistical Society. Series B (Statistical Methodology) 74(2), 245–266. URL: http://www.jstor.org/stable/41430939

Tukey, R. H. & Strassburg, C. P. (2000), ‘Human udp-glucuronosyltransferases: metabolism, expression, and disease’, Annual review of pharmacology and toxicology 40(1), 581–616.

